# Brain Signal Variability and Brain–Behavior Coupling in Adults with Dyslexia

**DOI:** 10.64898/2026.11.13.766529

**Authors:** Jie Luo, Arielle S. Keller, Zhichao Xia, Nikki Arrington, Lisa C. Krishnamurthy, Silvia Siu Yin Clement-Lam, Kenneth Pugh, Fumiko Hoeft

## Abstract

**Background:** Brain signal variability indexed by fMRI has emerged as a promising transdiagnostic biomarker for neurodevelopmental disorders, demonstrating better predictive utility than time-averaged metrics for diagnosis, yet it remains underexplored in adult dyslexia. The Neural Noise Hypothesis of dyslexia proposes that genetic risk disrupts cortical excitation/inhibition balance, elevating neural noise and affecting processes essential for reading. Here, we examine resting-state fMRI BOLD variability, a metric linked to neural noise, in adults with dyslexia and investigate its links with behavior.

**Methods:** In 30 adults with dyslexia (DY) and 76 typical readers (TR), we computed three complementary BOLD variability metrics: standard deviation (SD) and mean squared successive differences (MSSD) to quantify BOLD amplitude fluctuations, and dynamic regional homogeneity (dReHo) to quantify local temporal coordination. Voxel-wise group comparisons and partial least squares correlation analyses examined brain–behavior associations across a comprehensive behavioral battery spanning reading (e.g., word reading, phonological processing), math, oral language, working memory, and executive function domains.

**Results:** DY showed higher SD and MSSD in the left ventral occipitotemporal pathway and higher MSSD in the cerebellum, whereas TR showed higher dReHo in frontal and temporoparietal regions. SD/MSSD in visual, somatomotor, cerebellar, and subcortical networks significantly covaried with multi-domain behavioral profiles.

**Conclusions:** Adults with dyslexia showed localized, metric-specific differences in brain signal variability and brain–behavior covariance, indicating region-specific rather than global alterations. These results suggest that brain signal variability is a promising candidate task-free phenotype that could potentially index target engagement in interventions for adult dyslexia.

## Introduction

Dyslexia is a common neurodevelopmental disorder (prevalence 5–17.5%) (1,2) characterized by persistent reading difficulties arising from atypical neural processing, despite formal reading education (3). Adults with dyslexia continue to show deficits in reading fluency, pseudoword decoding, and spelling, alongside atypical function in reading-related networks, with lasting consequences for employment and quality of life (4,5,6). Yet, adults with dyslexia remain relatively understudied compared with developmental samples (7). Because brain function evolves across the lifespan, adults with dyslexia may show brain markers that differ from those identified in children. Thus, there is a strong need to identify brain markers of adults with dyslexia to inform mechanism-based, biologically grounded interventions for this population (8).

Moment-to-moment brain signal variability indexed by functional magnetic resonance imaging (fMRI) has recently emerged as a promising transdiagnostic biomarker of psychiatric symptoms that may apply across neurodevelopmental stages. Brain signal variability may better capture individual differences in diagnosis, prognosis, and treatment response than standard time-averaged measures (9,10). Importantly, a recent integrative framework has proposed that the moment-to-moment variability in fMRI signals may arise from variation in neuromodulatory systems that in turn lead to variation in excitation/inhibition (E/I) balance and microscale neural noise, which then manifest in macroscale brain signal variability and ultimately behavioral/symptom outcomes in psychiatric (e.g., major depressive disorder; MDD) and neurodevelopmental disorders (e.g., autism spectrum disorder; ASD) (11).

Despite this promising progress, it remains unknown whether adults with dyslexia are also characterized by variation in moment-to-moment brain signal variability. Among mechanistic accounts of dyslexia, the Neural Noise Hypothesis (NNH) proposes that genetic risk leads to disrupted cortical excitation–inhibition (E/I) balance and elevated neural noise, with this elevated noise thought to then contribute to reading difficulties (12). The NNH remains under active empirical investigation, with prior tests mostly focusing on E/I-related measures at the molecular or oscillatory level, such as magnetic resonance spectroscopy (MRS) and the aperiodic component of electroencephalograms (EEG) (13,14). Critically, E/I-driven changes in neural excitability alter spike timing and firing-rate regularity—microscale fluctuations that can be conceptualized as neural noise. These fluctuations propagate through local network synchronization, manifesting in moment-to-moment brain signal variability and, ultimately, individual differences in reading-related behavior (15,16,17). Resting-state fMRI (rs-fMRI) variability may therefore provide a macroscale proxy for E/I-related neural noise. Moment-to-moment brain signal variability has been linked to cognitive performance and functional brain organization (10,18,19), providing meaningful information beyond what can be gleaned from standard time-averaged measures. Brain signal variability has been linked to clinically relevant variation, including cognitive flexibility deficits in ASD and sustained attention deficits in attention-deficit/hyperactivity disorder (ADHD) (9,20).

Brain signal variability can be measured in multiple ways, each capturing distinct types of variation. Amplitude-based metrics include standard deviation (SD), which quantifies the magnitude of BOLD signal fluctuations over time, and mean squared successive difference (MSSD), which quantifies the average change in BOLD signal across consecutive timepoints. These amplitude-based measures have been linked to laminar cortical architecture in granular layer IV, ionotropic receptor density, E/I balance, and metabolic activity (21). Cross-species evidence further suggests that low-frequency BOLD fluctuations may reflect underlying neural activity, consistent with a physiologically plausible hemodynamic delay (22,23). Another class of metrics quantifies local coupling dynamics. For example, dynamic regional homogeneity (dReHo) quantifies moment-to-moment variability in local synchrony between a voxel and its neighbors, indexing short-range cortical coordination. Variability in coupling dynamics has been linked to infragranular layers V–VI, metabotropic receptor density, and large-scale connectivity gradients (21,24,25). A recent study found that anesthesia consistently reduces the temporal autocorrelation of local BOLD signals across humans, nonhuman primates, and mice, with these changes covarying with cortical expression of excitatory/inhibitory neurotransmission genes (26). Together, amplitude-based and coupling-dynamics metrics index complementary, biologically grounded aspects of brain signal variability.

Prior fMRI studies of adults with dyslexia have primarily relied on time-averaged measures that do not capture the temporal dynamics central to E/I-driven neural noise (27) and fall short of explaining how local dynamics give rise to distributed activity—a link that requires investigating moment-to-moment BOLD variability. Although recent work has examined task-based, trial-by-trial BOLD fluctuations related to reading performance and genetic risk (28,29), it remains unknown whether intrinsic resting-state brain signal variability differs between adults with and without dyslexia. Several key questions remain unresolved. First, are differences in variability detectable at rest, independent of task demands? Second, are such differences spatially localized to reading-relevant regions, such as the left ventral occipitotemporal cortex (30) and dorsal temporoparietal pathway (4), or broadly distributed? Third, does brain variability preferentially relate to reading and language abilities or to multi-domain cognitive skills?

Inspired by the NNH framework, the present study addresses these questions by using rs-fMRI variability as a task-free marker of dyslexia. We use amplitude-based and coupling-dynamics metrics to test whether dyslexia-related variability differences are 1) detectable at the group level, by comparing SD, MSSD, and dReHo between adults with dyslexia (DY) and typical readers (TR); 2) spatially localized, by testing whether effects fall within predefined reading-network masks or are broadly distributed; and 3) behaviorally specific, by using multivariate partial least squares correlation (PLS-C) to examine whether brain variability covaries with reading/language or multi-domain behaviors, and whether this brain-behavior covariance differs between DY and TR. Together, these questions investigate whether adult dyslexia is characterized by disrupted moment-to-moment rs-fMRI brain signal variability, and whether such disruptions explain differences in reading-specific or multi-domain behaviors.

## Methods and Materials

### Participants

Participants with available data in this dataset (*N*=150, ages 18.08-39.31 years old) were recruited and tested at two sites: the Center for Advanced Brain Imaging (CABI, Georgia State University, Atlanta, GA; *n*=34) and the Brain Imaging Research Center (BIRC, University of Connecticut, Storrs, CT; *n*=116). 44 participants were excluded for incomplete data (missing rs-fMRI and/or behavioral assessment data; n=41) or failed quality control (n=3; see Data Preprocessing), yielding a final sample of 106. DY (n = 30) had Test of Word Reading Efficiency–2 Phonemic Decoding Efficiency (PDE) < 90 and a self-reported history of childhood reading difficulties assessed through the Adult Reading History Questionnaire, ARHQ ≥ 0.40 (31). The typical-reader group (TR; *n* = 76) had typical reading (PDE ≥ 90) and no documented history (ARHQ < 0.40). All participants were neurologically healthy, right-handed, native English-speaking young adults with normal or corrected vision and hearing and a Full-Scale IQ ≥ 85. Groups were matched in age (DY: 22.24 ± 3.97; TR: 23.18 ± 5.35 years) and sex (DY: 9/21; TR: 29/47 male/female). All study procedures were approved by the UConn IRB #HR18-112GSU and conducted in accordance with the Declaration of Helsinki, and all participants provided written informed consent.

### Behavior Assessment and Neuroimaging Data

Participants completed a battery of standardized assessments of reading (word and pseudoword reading, fluency, spelling, phonological processing), oral language, math, working memory, executive function, and psychological functioning (full list in Supplement). MRI data were acquired on a 3 Tesla Siemens Prisma scanner. T1-weighted anatomical images were obtained using a 3D MPRAGE sequence (repetition time = 1,900 ms; echo time = 2.52 ms; FOV = 224 mm; 1 mm isotropic; 176 sagittal slices). Eyes-open, video-based (Inscapes) (32) rest naturalistic-viewing fMRI used a multi-echo sequence (repetition time = 1,630 ms; echo times = 16.00/37.79/59.58 ms; flip angle = 67°; FOV = 210 mm; matrix = 84 × 84; 2.5 mm isotropic; 54 axial slices; 377 volumes; 10 min 16 s). The multi-echo design improves temporal signal-to-noise ratio (tSNR) by leveraging TE-dependent BOLD signal separation (33). Reversed phase-encode (AP/PA) spin-echo field maps were also acquired for susceptibility distortion correction.

### Data Preprocessing

Because BOLD brain variability is sensitive to artifacts, preprocessing extended beyond standard steps to include within-subject ICA denoising, white matter (WM), cerebrospinal fluid (CSF), and motion-parameter regression, and block normalization, which reduce voxel-level variance and improve statistical prediction (34,35,36). Preprocessing was performed in AFNI v23.0.00 (37) and included quality inspection, removal of the first five volumes, slice-timing and AP/PA distortion correction, EPI–T1 coregistration, motion correction estimated in native space, combined with the subject-specific nonlinear warp into MNI152 space in a single resampling step, multi-echo combination and ME-ICA denoising via tedana (33; performed in MNI space after normalization to enable atlas-based component classification), scaling, WM/CSF nuisance regression, detrending, censoring of high head motion/outlier volumes, band-pass filtering (0.01– 0.1 Hz), and 4 mm FWHM smoothing. Participants were excluded (n=3, of the 44 above) at quality-control steps for poor spatial alignment by visual inspection, signal anomalies (e.g., signal dropout or ghosting), excessive head motion (mean framewise displacement (*mean*FD; the average head displacement) >0.2mm), or insufficient TE-dependent (BOLD-like) ICA components (indicating poor signal separation in multi-echo ICA decomposition).

### Brain Variability Metrics Calculation

After preprocessing, SD, MSSD, and dReHo of brain variability were computed in AFNI. For both SD and MSSD, Xt represents the BOLD signal at time t, i.e., the signal value of a given voxel or region at each time point, and T is the total number of time points (372). For dReHo, (*M*) in each sliding window M is computed from the windowed time series (for = 1,…, where is the window length) by quantifying voxel-wise concordance with neighboring voxels using Kendall’s coefficient of concordance. In line with prior work (38), the main dReHo analysis used a 62-volume window with a 5-volume step (dReHo(w62s5)), given the 0.01 Hz band-pass cutoff; alternative configurations (w62s1, w30s5, w30s1) were tested for robustness (Supplemental Methods).

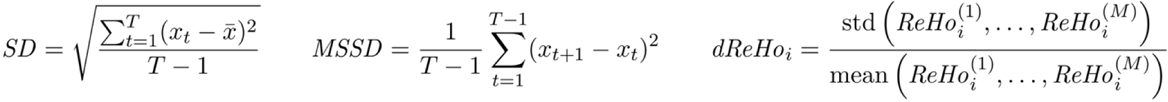

### Statistical Analyses

#### Group Differences in Brain Variability

Voxel-wise general linear models (GLMs) compared TR and DY on each metric at the whole-brain level and within two reading-network masks: a Neurosynth meta-analytic mask (term “Reading”; 39; broad, data-driven coverage) and a literature-based mask mapped onto the Schaefer-400 atlas, comprising all 153 parcels containing at least one reading-related peak coordinate (40-41; targeted parcellation-level coverage, see Supplemental Methods). All models included covariates for age, sex, scanning site (group distribution across sites was comparable), meanFD (42), and Barkley ADHD current total score (43). Given the co-occurrence of dyslexia with ADHD and their independent influence on brain variability (9,20), ADHD symptoms were included as covariates rather than used as an exclusion criterion to preserve ecological validity. Multiple comparisons were FWE-corrected (voxel *p* < .005 or .01; cluster *p*FWE < .05) using AFNI’s 3dClustSim with residual-based autocorrelation function (ACF) parameters (44). For validation and interpretation, we also computed whole-brain mean variability group comparisons for each metric, inter-metric correlations, tSNR and *mean*FD validation to rule out data quality confounds (45,46), and dReHo window-parameter sensitivity tests (Supplemental Methods).

#### Multivariate Brain–Behavior Associations and Group Differences Through Partial Least Squares Correlation (PLS-C) Analyses

PLS-C was implemented in Python (v3.12.7) to identify multivariate associations between whole-brain voxel-wise variability metrics and behavioral measures. PLS-C is typically more robust for high-dimensional data (features > observations) and strongly dependent predictors (e.g., spatial autocorrelation in neuroimaging maps or correlated behavioral measures), making it well-suited for voxel-wise neuroimaging applications. Two models per metric differed only in the behavioral feature set: 1) a multi-domain model (*n* = 22 variables assessing reading, language, mathematics, executive function, and working memory behavior); and 2) a reading/language-specific model (*n* = 13 variables). Details are provided in Supplemental Methods.

PLS-C followed established practices (47,48,49,50). Before PLS-C analysis, behavioral matrices were rank-based inverse normal transformed (ranks each variable and maps it to a normal distribution, providing robustness to outliers without assuming parametric distributions), brain and behavioral matrices were then covariate-regressed (age, sex, *mean*FD, scanning site, and Barkley ADHD Current Total score), and z-scored. PLS-C was then applied to the combined sample (*N* = 106) by computing the cross-covariance matrix R = XTY/(n−1) between brain data X (n × p voxels) and behavior Y (n × q), then decomposing R = USVT via singular value decomposition (SVD), yielding latent components (LCs) with associated brain and behavior saliences (weight vectors in U and V, respectively, S contains singular values). Brain scores (T = XU) and behavioral scores (S = YV) were computed by projecting the preprocessed data onto the respective saliences. LC significance used within-group permutation (2,000); feature reliability used bootstrap resampling (1,000 iterations; |Bootstrap Ratios (BSR)| > 3.0 reliable). Significant voxels were mapped to the Schaefer 400/7-network (41), Tian subcortical (51), and SUIT cerebellar (52) atlases. Full details in Supplemental Methods.

Two complementary tests characterized group differences along significant LCs. Δscore tested whether groups differed in LC expression: brain scores (T) and behavioral scores (S) derived from the full-sample weights were compared across groups, with significant difference indicating that DY and TR occupy distinct positions along the same latent brain–behavior axis (49,53). Δr quantified brain–behavior coupling strength: within-group brain–behavior correlations on the same LC (*r*TR, *r*DY) generated Δr = *r*TR – *r*DY, with significance indicating differences in coupling coherence, a stricter test than Δscore (48,53). Sensitivity analyses for validation and interpretation included 5-fold cross-validation, spin-based permutation testing to account for spatial autocorrelation (54,55) and dReHo sliding window-length and steps sensitivity tests (Supplemental Methods).

## Results

### Group Differences in Resting-state fMRI Brain Variability Between Adults with and without Dyslexia

To examine group differences in BOLD variability, we compared TR and DY voxel-wise across all three metrics. At the whole-brain level (voxel-wise *p* < .005, cluster-level *p*FWE < .05), group differences emerged for all three metrics (Figure 1; positive t-values indicate TR > DY, negative t-values indicate DY > TR). The DY group showed higher SD and MSSD in the left ventral occipitotemporal reading pathway (inferior temporal gyrus) and higher MSSD in the left cerebellum, while TR showed higher dReHo in the right superior frontal and middle temporal gyri and the left dorsal temporoparietal reading pathway (supramarginal gyrus). Within reading-network masks (voxel-wise *p* < .01, cluster-level *p*FWE < .05), only dReHo differed between groups. In the literature-based mask, TR showed higher dReHo than DY in frontal regions (left superior medial frontal cortex, right middle frontal gyrus), the left ventral occipitotemporal pathway (fusiform and middle occipital gyri), and the left supramarginal gyrus (Figure 2); in the Neurosynth-derived mask, TR showed higher dReHo in the left fusiform gyrus (Figure 3). Coordinates and cluster statistics are reported in Table 1.

**Figure 1.**
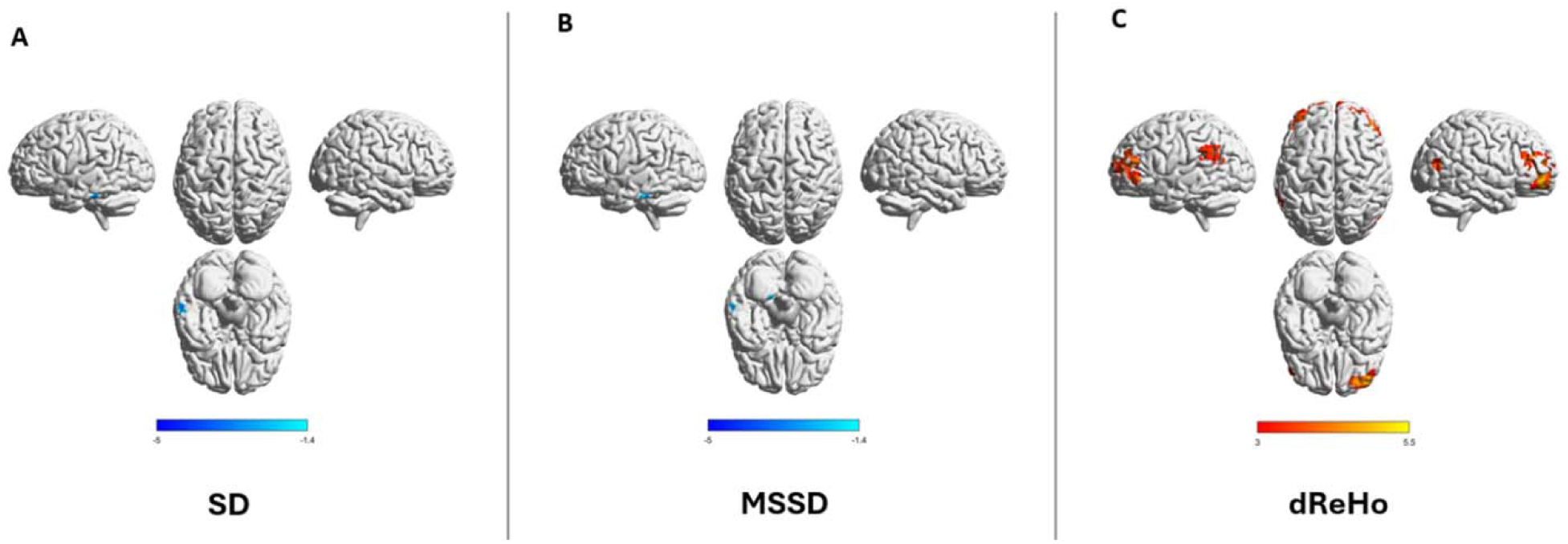
Whole-brain group differences in resting-state fMRI signal variability between typical readers (TR) and dyslexia (DY). (A) Standard Deviation (SD), (B) Mean Squared Successive Difference (MSSD), (C) Dynamic Regional Homogeneity (dReHo). The color bar represents the t-statistic of the group contrast. For SD and MSSD, blue color (negative t-values) indicates lower SD and MSSD in the TR group relative to the DY group. For dReHo, red-yellow color (positive t-values) indicates higher dReHo in the TR group relative to the DY group. All maps were thresholded at a voxel-wise *p* < .005. Cluster-level significance was determined to be *p*_FWE_ < .05 using AFNI’s 3dClustSim, with the minimum cluster extent estimated from residual-based spatial autocorrelation parameters (ACF) to control for multiple comparisons. Results are displayed in Montreal Neurological Institute (MNI) coordinates.

**Figure 2.**
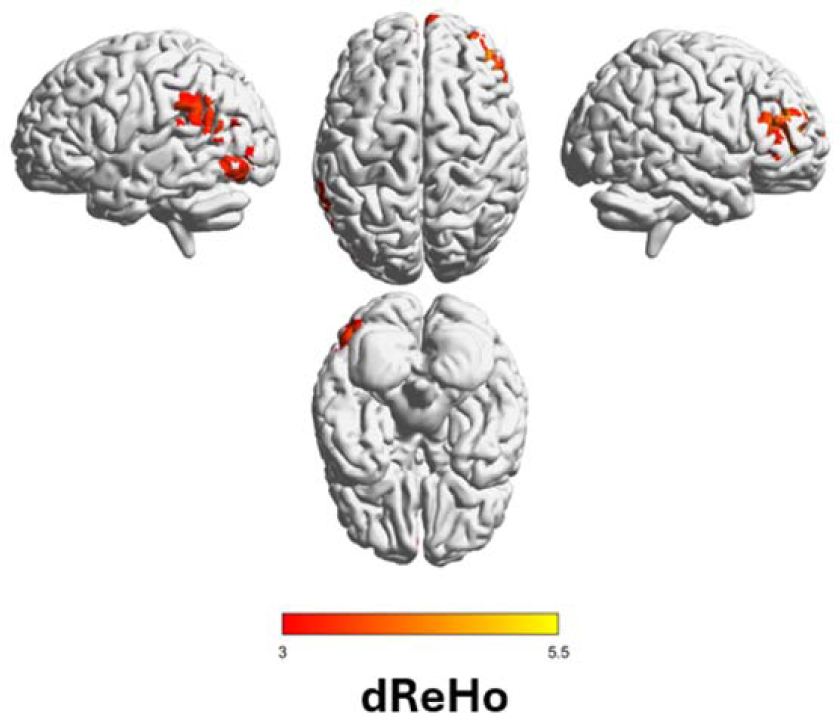
Reading-network (defined from previous literature) group differences in resting-state fMRI signal variability between typical readers (TR) and dyslexia (DY). Dynamic Regional Homogeneity (dReHo). The color bar represents the t-statistic of the group contrast. For dReHo, red-yellow color (positive t-values) indicate higher dReHo in the TR group relative to the DY group. All maps were thresholded at a voxel-wise *p* < .01. Cluster-level significance was determined to be *p*_FWE_ < .05 using AFNI’s 3dClustSim, with the minimum cluster extent estimated from residual-based spatial autocorrelation parameters (ACF) to control for multiple comparisons. Results are displayed in Montreal Neurological Institute (MNI) coordinates.

**Figure 3.**
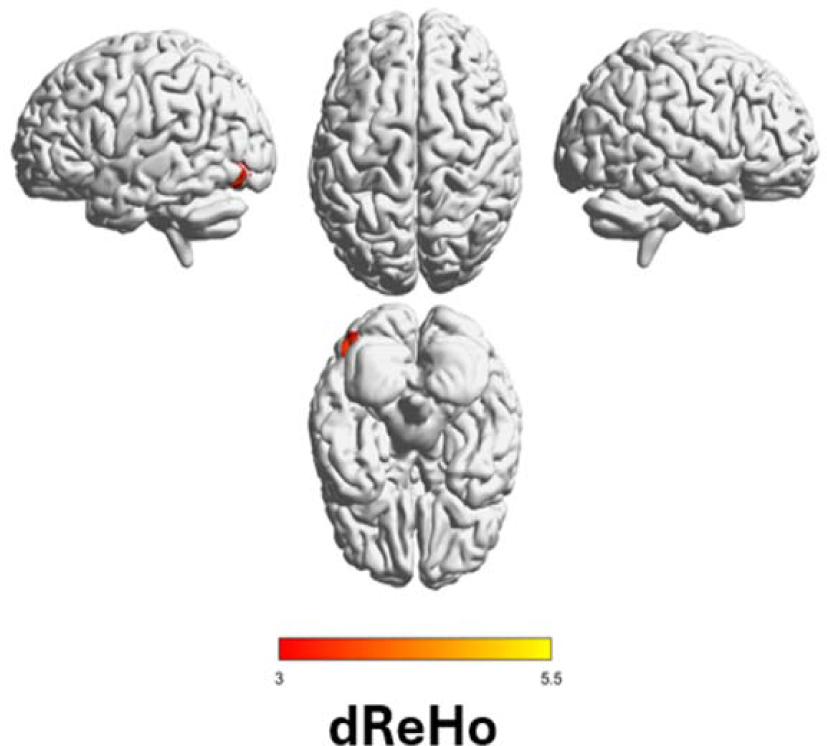
Reading-network (defined from Neurosynth) group differences in resting-state fMRI signal variability between typical readers (TR) and dyslexia (DY). Dynamic Regional Homogeneity (dReHo). The color bar represents the t-statistic of the group contrast. For dReHo, red-yellow color (positive t-values) indicate higher dReHo in the TR group relative to the DY group. All maps were thresholded at a voxel-wise *p* < .01. Cluster-level significance was determined to be *p*_FWE_ < .05 using AFNI’s 3dClustSim, with the minimum cluster extent estimated from residual-based spatial autocorrelation parameters (ACF) to control for multiple comparisons. Results are displayed in Montreal Neurological Institute (MNI) coordinates.

**Table 1.** Significant clusters from voxel-wise group comparisons (TR vs. DY) at the whole-brain level and within reading-network masks.

| Analysis | Metric | Hemisphere | Region | x | y | z | Cluster size (voxels) | Peak t |
|---|---|---|---|---|---|---|---|---|
| **Whole-brain** | SD | Left | ITG | −60 | −33 | −26 | 66 | −3.8 |
|  |  | Left | ITG | −58 | −32 | −25 | 57 | −3.9 |
|  | MSSD | Left | Cerebellum | −13 | −47 | −53 | 48 | −4.1 |
|  | dReHo | Right | SFG | 18 | 52 | 8 | 945 | 5.6 |
|  |  | Right | MTG | 45 | −73 | 13 | 78 | 4.3 |
|  |  | Left | SMG | −63 | −53 | 26 | 72 | 3.8 |
|  |  | Right | MFG | 42 | 44 | 15 | 289 | 5.6 |
| **Reading-network (literature)** | dReHo | Left | Superior medial frontal gyrus | −2 | 58 | 16 | 134 | 4.2 |
|  |  | Left | FFG | −46 | −76 | −9 | 60 | 3.6 |
|  |  | Left | MOG | −42 | −78 | 9 | 55 | 3.7 |
|  |  | Left | SMG | −62 | −53 | 25 | 87 | 3.8 |
| **Reading-network (Neurosynth)** | dReHo | Left | FFG | −42 | −80 | −11 | 29 | 3.6 |

Note. ITG = Inferior temporal gyrus; SFG = Superior frontal gyrus; MTG = Middle temporal gyrus; SMG = Supramarginal gyrus; MFG = Middle frontal gyrus; FFG = Fusiform gyrus; MOG = Middle occipital gyrus. Whole-brain analyses thresholded at voxel-wise *p* < .005, cluster *p*FWE < .05; reading-network analyses at voxel-wise *p* < .01, cluster *p*FWE < .05. Coordinates in MNI space.

### Multivariate Resting-state fMRI Brain Variability is Associated with Both Multi-domain and Reading/Language-specific Skills

#### Latent Component Structure Among Typical Readers (TR) and Dyslexia (DY)

To characterize latent brain–behavior structure across the full behavioral domain, we applied PLS-C with the multi-domain feature set (22 variables). For SD, PLS-C identified a significant LC1 (*p*_permuted_ = .006, 43.55% covariance; Figure 4A); 16 behavioral variables showed reliable contributions (|BSR| > 3; Figure 4B). Brain saliences (voxel-level weights reflecting each region’s contribution to the LC) were localized predominantly to somatomotor, visual, cerebellar, and subcortical networks (Figure 4C-D; voxel counts and |BSR| in Table S1). For MSSD, PLS-C identified a significant LC1 (*p*_permuted_ = .002, 43.44% covariance; Figure 5A); 16 behavioral variables contributed reliably (Figure 5B). Brain saliences showed the same network distribution as SD (Figure 5C–D; Table S2). For dReHo, no LC reached significance (LC1: *p*_permuted_ = .131; Figure S17).

**Figure 4.**
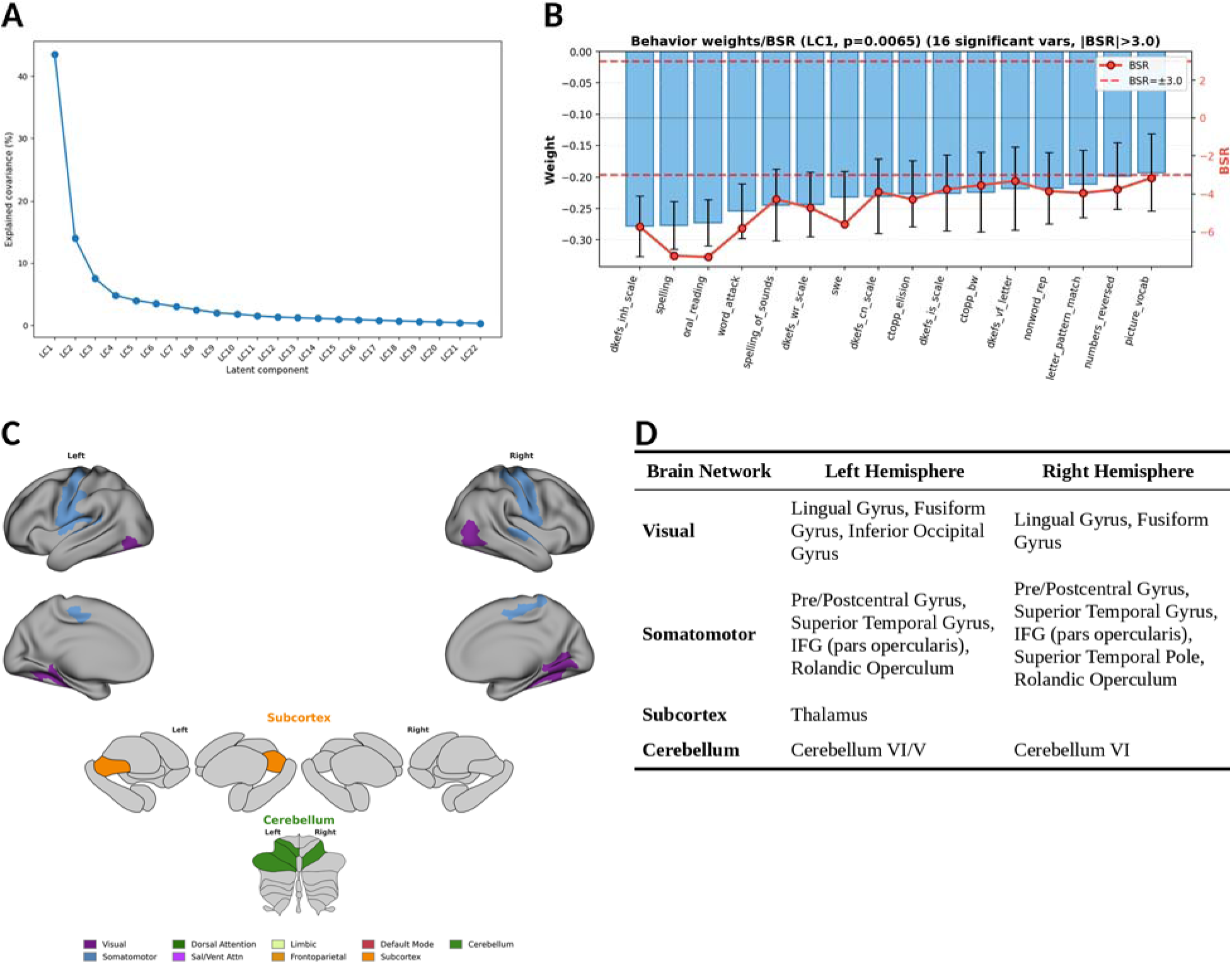
Standard deviation of resting-state fMRI BOLD signal (SD) and behavioral associations in the multi-domain model (all behavioral measures, including reading, language, math, working memory, and executive function) revealed by Partial Least Squares Correlation (PLS-C) analysis. (A) Variance explained by latent components. LC1 accounts for the largest shared variance between SD and behavioral measures (*p* < .05, permutation test). (B) Behavioral weights with bootstrap uncertainty (black error bars) and bootstrap ratios (BSR, red line with 95% CI) for significant behavioral variables (|BSR| > 3.0). Dashed lines: ±3 BSR threshold. Behavioral variable abbreviations: D-KEFS = Delis–Kaplan Executive Function System; CTOPP-2 = Comprehensive Test of Phonological Processing–2nd Ed. Individual variable names and full descriptions are listed in Supplemental Methods (Behavioral Measures). (C) Spatial distribution of significant voxel-level BSR values across the brain. Cortical regions are mapped onto Yeo 7-network parcellation; subcortical and cerebellar regions are displayed using the subcortex visualization toolbox, shown in four views (left lateral, left medial, right lateral, right medial) so that both laterally and medially located nuclei are visible (ROI-level mapping; only regions with ≥10 significant voxels are projected onto the template; regions below this threshold are shown at actual voxel locations). (D) Brain regions with significant BSR values (|BSR| > 3.0) organized by network and hemisphere. Regions are grouped according to the Yeo 7-network parcellation (cortical), Tian atlas (subcortical), and SUIT atlas (cerebellar). For visualization clarity, the top regions ranked by number of significant voxels are displayed; regions with fewer significant voxels are omitted. 2000 permutations for LC significance (*p* < .05), 1000 bootstrap iterations for BSR (|BSR| > 3.0). N = 106 (Typical Readers: 76, Dyslexia: 30). Brain parcellation: multi-atlas approach (Schaefer 400-parcel cortical, Tian subcortical, SUIT cerebellar atlases). Network abbreviations (Panel C abbreviation = Panel D full name): SomMot = Somatomotor, Vis = Visual, Cerebellum = Cerebellar, Subcortex = Subcortical. IFG = inferior frontal gyrus.

**Figure 5.**
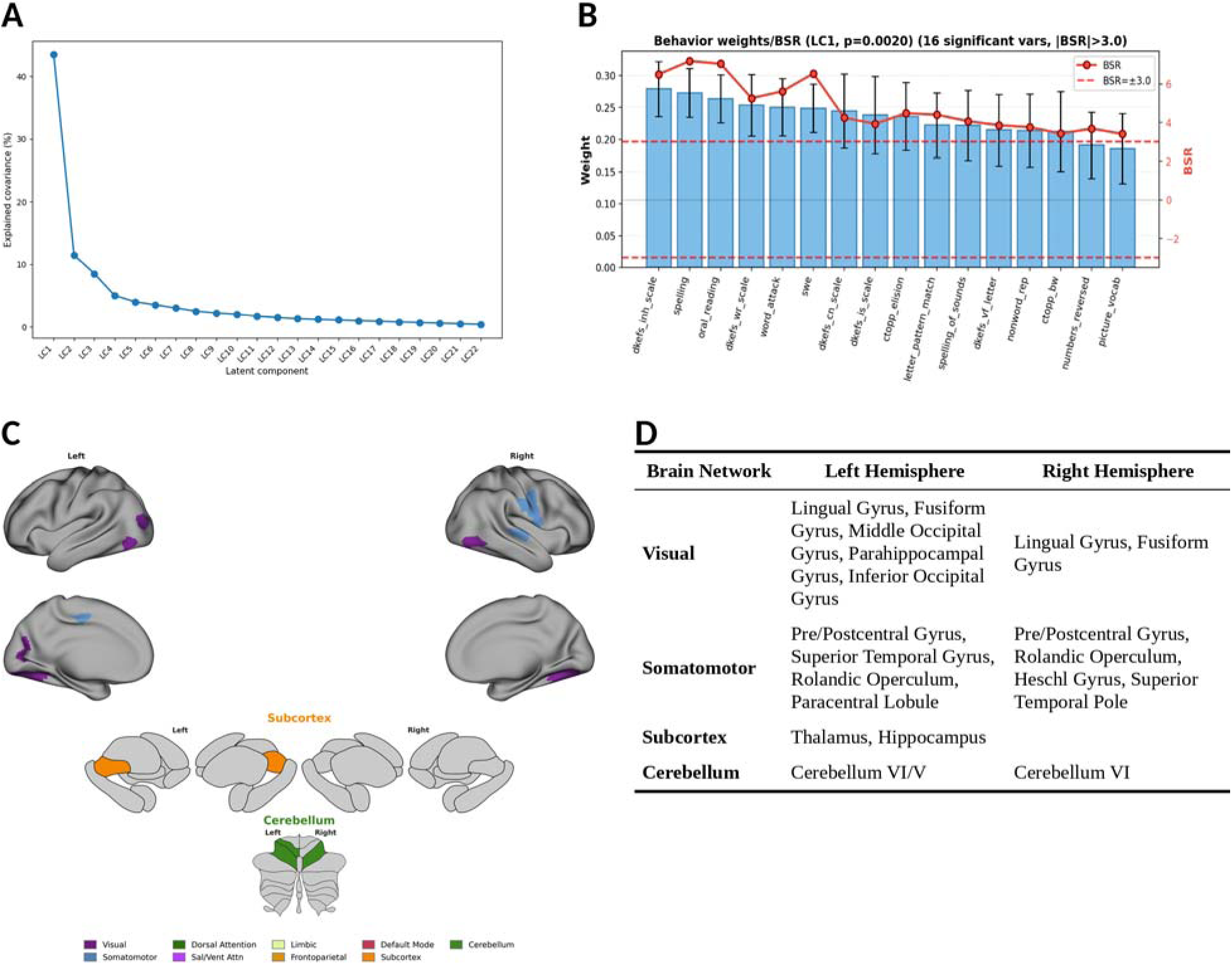
Mean Squared Successive Difference of resting-state fMRI BOLD signal (MSSD) and behavioral associations in the multi-domain model (all behavioral measures, including reading, language, math, working memory, and executive function) revealed by Partial Least Squares Correlation (PLS-C) analysis. (A) Variance explained by latent components. LC1 accounts for the largest shared variance between MSSD and behavioral measures (*p* < .05, permutation test). (B) Behavioral weights with bootstrap uncertainty (black error bars) and bootstrap ratios (BSR, red line with 95% CI) for significant behavioral variables (|BSR| > 3.0). Dashed lines: ±3 BSR threshold. Behavioral variable abbreviations: D-KEFS = Delis–Kaplan Executive Function System; CTOPP-2 = Comprehensive Test of Phonological Processing–2nd Ed. Individual variable names and full descriptions are listed in Supplemental Methods (Behavioral Measures) (C) Spatial distribution of significant voxel-level BSR values across the brain. Cortical regions are mapped onto Yeo 7-network parcellation; subcortical and cerebellar regions are displayed using the subcortex visualization toolbox, shown in four views (left lateral, left medial, right lateral, right medial) so that both laterally and medially located nuclei are visible (ROI-level mapping; only regions with ≥10 significant voxels are projected onto the template; regions below this threshold are shown at actual voxel locations). (D) Brain regions with significant BSR values (|BSR| > 3.0) organized by network and hemisphere. Regions are grouped according to the Yeo 7-network parcellation (cortical), Tian atlas (subcortical), and SUIT atlas (cerebellar). For visualization clarity, the top regions ranked by number of significant voxels are displayed; regions with fewer significant voxels are omitted. 2000 permutations for LC significance (*p* < .05), 1000 bootstrap iterations for BSR (|BSR| > 3.0). N = 106 (Typical Readers: 76, Dyslexia: 30). Brain parcellation: multi-atlas approach (Schaefer 400-parcel cortical, Tian subcortical, SUIT cerebellar atlases). Network abbreviations (Panel C abbreviation = Panel D full name): SomMot = Somatomotor, Vis = Visual, Cerebellum = Cerebellar, Subcortex = Subcortical.

To test whether these patterns exist when restricting to reading- and language-relevant behavior, we repeated PLS-C with the reading/language-specific feature set (13 variables). For SD, PLS-C identified a significant LC1 (*p*_permuted_ = .019, 50.02% covariance; Figure S15A); 10 behavioral variables contributed reliably (Figure S15B). Brain saliences localized to somatomotor, visual, cerebellar, and subcortical networks (Figure S15C–D; Table S3). For MSSD, PLS-C identified a significant LC1 (*p*_permuted_ = .009, 49.41% covariance; Figure S16A); 10 behavioral variables contributed reliably (Figure S16B). Brain saliences showed the same network distribution (Figure S16C–D; Table S4). For dReHo, no LC reached significance (LC1: *p*_permuted_ = .220).

#### Measures Capturing Variability in the Amplitude of Local Fluctuations Show Multi-domain and Reading-specific Patterns of Brain-Behavior Covariance that Differentiate Adults with and without Dyslexia

To test whether amplitude-based variability differentiates groups in their brain–behavior associations, we applied group-comparison PLS-C to SD and MSSD. In the multi-domain model, both SD-LC1 and MSSD-LC1 showed significant group differences in LC expression (SD: *t*(104) = −4.97/−9.61, *d* = −1.27/−2.07; MSSD: *t*(104) = 5.24/9.60, *d* = 1.44/2.07; all *p* < .001; Figures 6A, 7A). Brain–behavior coupling strength did not differ between groups for either metric (SD: *r*TR = 0.400, *r*DY = 0.638, *p*_permuted_ = .084; MSSD: *r*TR = 0.489, *r*DY = 0.660, *p*_permuted_ = .138; Figures 6B–C, 7B–C). For SD, one additional LC (LC10) showed a significant Δr but had non-significant brain–behavior covariance and was not interpreted further.

**Figure 6.**
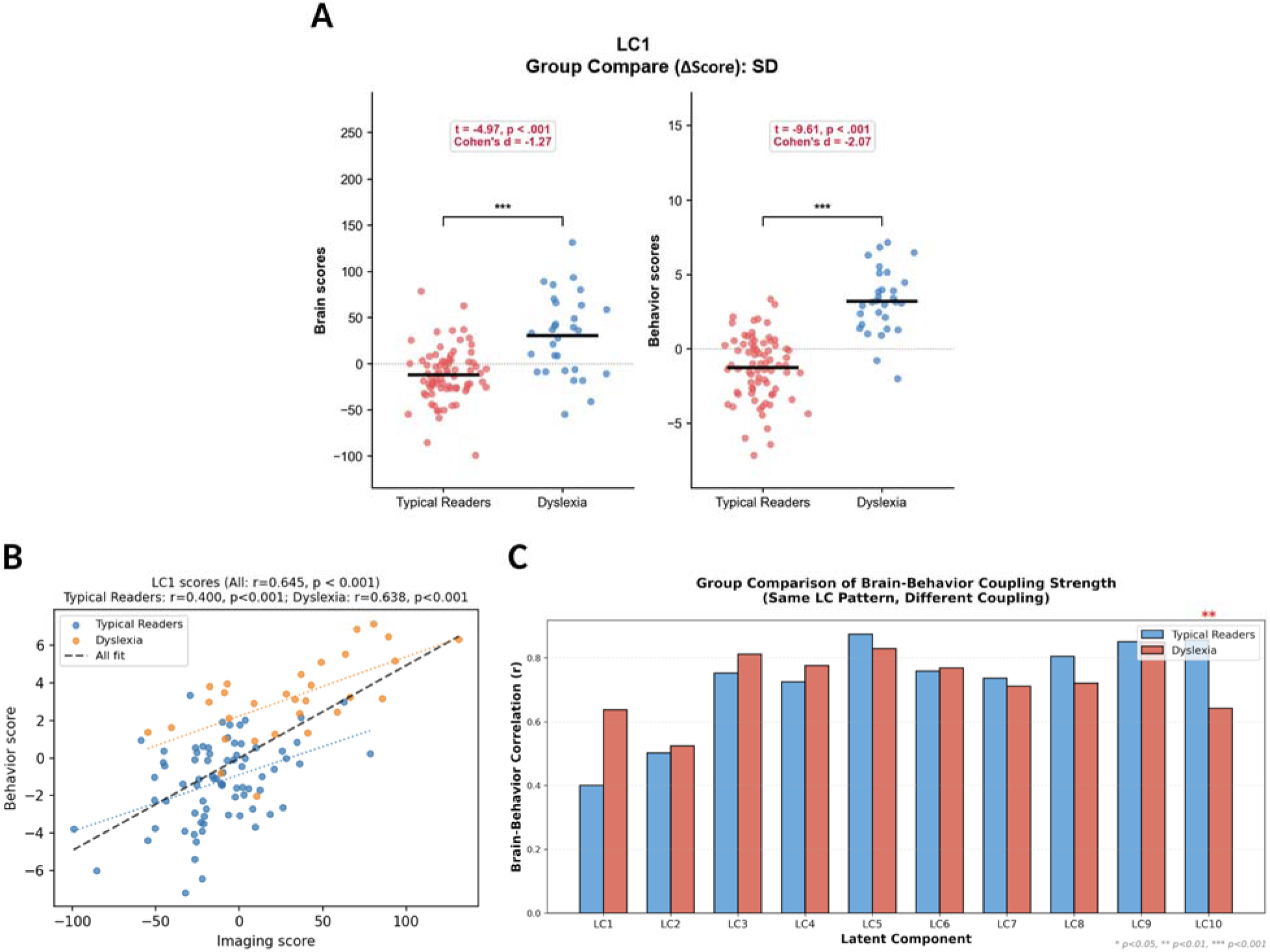
Group differences in brain-behavior associations revealed by Partial Least Squares Correlation (PLS-C) analysis of resting-state fMRI BOLD signal standard deviation (SD) and the multi-domain behavioral model between typical readers (TR) and dyslexia (DY). (A) Group comparisons of LC1 brain and behavioral scores. Red: TR; Blue: DY. (B) Brain-behavior coupling strength (*r*) in LC1. Brain scores (x-axis) vs. behavioral scores (y-axis). Blue: TR; Orange: DY. (C) Brain-behavior coupling strength (Δr = rTR − rDY) across LC1-LC10. Significant differences mean brain-behavior coupling differed between groups on the same shared LC (\**p* < .05, \*\**p* < .01, \*\*\**p* < .001). Blue: TR; Red: DY. Statistical testing: Δr significance assessed via label permutation (2,000 iterations). TR: n = 76, DY: n = 30.

**Figure 7.**
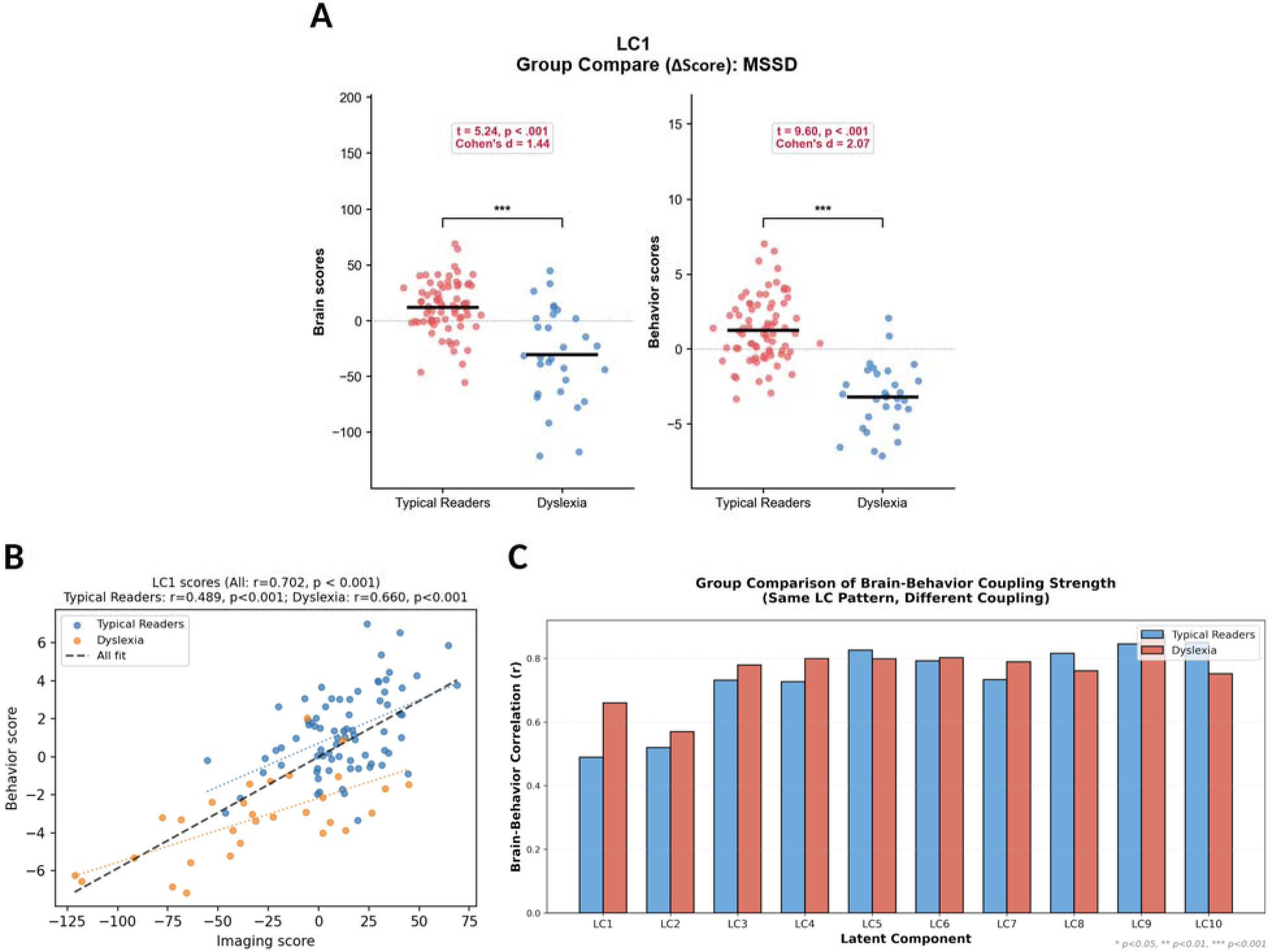
Group differences in brain-behavior associations revealed by Partial Least Squares Correlation (PLS-C) analysis of resting-state fMRI BOLD signal Mean Squared Successive Difference (MSSD) and the multi-domain behavioral model between typical readers (TR) and dyslexia (DY). (A) Group comparisons of LC1 brain and behavioral scores. Red: TR; Blue: DY. (B) Brain-behavior coupling strength (*r*) in LC1. Brain scores (x-axis) vs. behavioral scores (y-axis). Blue: TR; Orange: DY. (C) Brain-behavior coupling strength (Δr = rTR − rDY) across LC1-LC10. Significant differences mean brain-behavior coupling differed between groups on the same shared LC (\**p* < .05, \*\**p* < .01, \*\*\**p* < .001). Blue: TR; Red: DY. Statistical testing: Δr significance assessed via label permutation (2,000 iterations). TR: n = 76, DY: n = 30.

Results were similar in the reading/language-specific model: SD-LC1 and MSSD-LC1 again showed significant group differences in LC expression (SD: *t*(104) = −4.46/−8.40, *d* = −1.07/−1.72; MSSD: *t*(104) = 4.73/8.44, *d* = 1.23/1.74; all *p* < .001; Figures S18A, S19A), and brain–behavior coupling strength did not differ between groups (SD: *r*TR = 0.434, *r*DY = 0.600, *p*_permuted_ = .251; MSSD: *r*TR = 0.493, *r*DY = 0.637, *p*_permuted_ = .263; Figures S18B–C, S19B–C).

#### Variability in Local Coherence Measured by dReHo Shows Multi-domain Brain-Behavior Covariance that Differentiates Adults with and without Dyslexia

Although dReHo did not yield a significant LC in the combined-sample analysis, which tests for a covariance pattern shared across all participants, group-comparison PLS-C, which tests whether DY and TR differ in expression of or coupling along latent dimensions, identified a significant LC1, indicating a group-dependent rather than sample-wide pattern (*p*_permuted_ = .009, 49.41% covariance); 13 behavioral variables contributed reliably (Figure 8A). Brain saliences localized to somatomotor, visual, cerebellar, and subcortical networks (Figure 8B–C). Groups differed significantly in LC expression on brain (*t*(104) = −5.17, d = −1.33) and behavioral scores (*t*(104) = −9.54, *d* = −2.08; both *p* < .001; Figure 9A). Critically, dReHo also showed a significant brain–behavior coupling difference between TR and DY (*r*TR = 0.328 vs. *r*DY = 0.681; Δr = −0.353, *p*_permuted_ = .017; Figure 9B–C).

**Figure 8.**
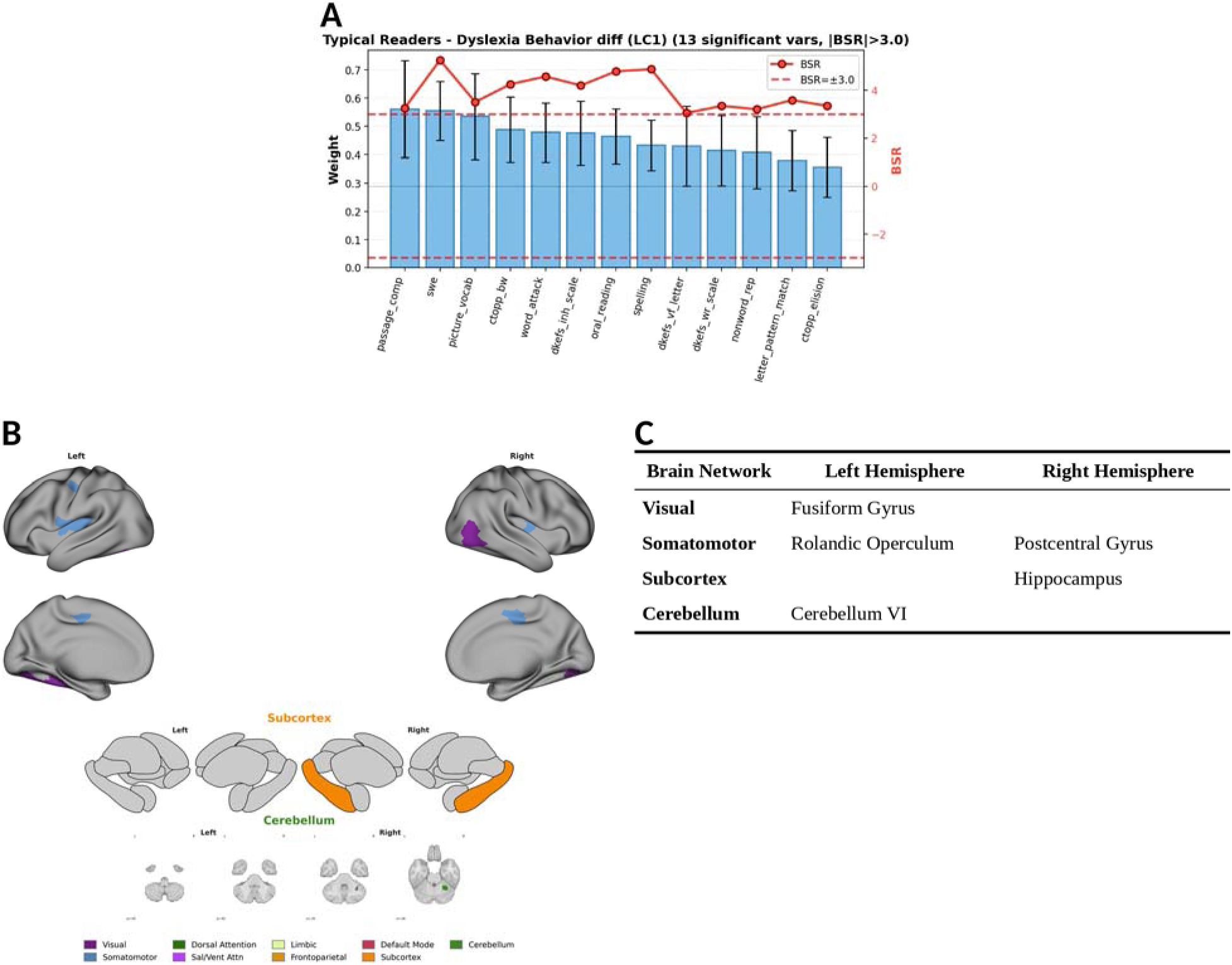
Dynamic Regional Homogeneity (dReHo) and behavioral associations in the multi-domain model revealed by group-comparison Partial Least Squares Correlation (PLS-C) analysis. (A) Behavioral weights with bootstrap uncertainty (black error bars) and bootstrap ratios (BSR, red line with 95% CI) for significant behavioral variables (|BSR| > 3.0). Dashed lines: ±3 BSR threshold. Behavioral variable abbreviations: D-KEFS = Delis–Kaplan Executive Function System; CTOPP-2 = Comprehensive Test of Phonological Processing–2nd Ed. Individual variable names and full descriptions are listed in Supplemental Methods (Behavioral Measures). (B) Spatial distribution of suprathreshold voxel-level BSR values across the brain. Cortical regions are mapped onto Yeo 7-network parcellation; subcortical regions are displayed using the subcortex visualization toolbox, shown in four views (left lateral, left medial, right lateral, right medial) so that both laterally and medially located nuclei are visible (ROI-level mapping; only regions with ≥10 significant voxels are projected onto the template; regions below this threshold are shown at actual voxel locations). Cerebellar regions are displayed at actual voxel locations, as no cerebellar ROI contained ≥10 suprathreshold voxels. (C) Brain regions with significant BSR values (|BSR| > 3.0) organized by network and hemisphere. Regions are grouped according to the Yeo 7-network parcellation (cortical), Tian atlas (subcortical), and SUIT atlas (cerebellar). 2000 permutations for LC significance (*p* < .05), 1000 bootstrap iterations for BSR (|BSR| > 3.0). N = 106 (Typical Readers: 76, Dyslexia: 30). Brain parcellation: multi-atlas approach (Schaefer 400-parcel cortical, Tian subcortical, SUIT cerebellar atlases). Network abbreviations (Panel B abbreviation = Panel C full name): SomMot = Somatomotor, Vis = Visual, Cerebellum = Cerebellar, Subcortex = Subcortical.

**Figure 9.**
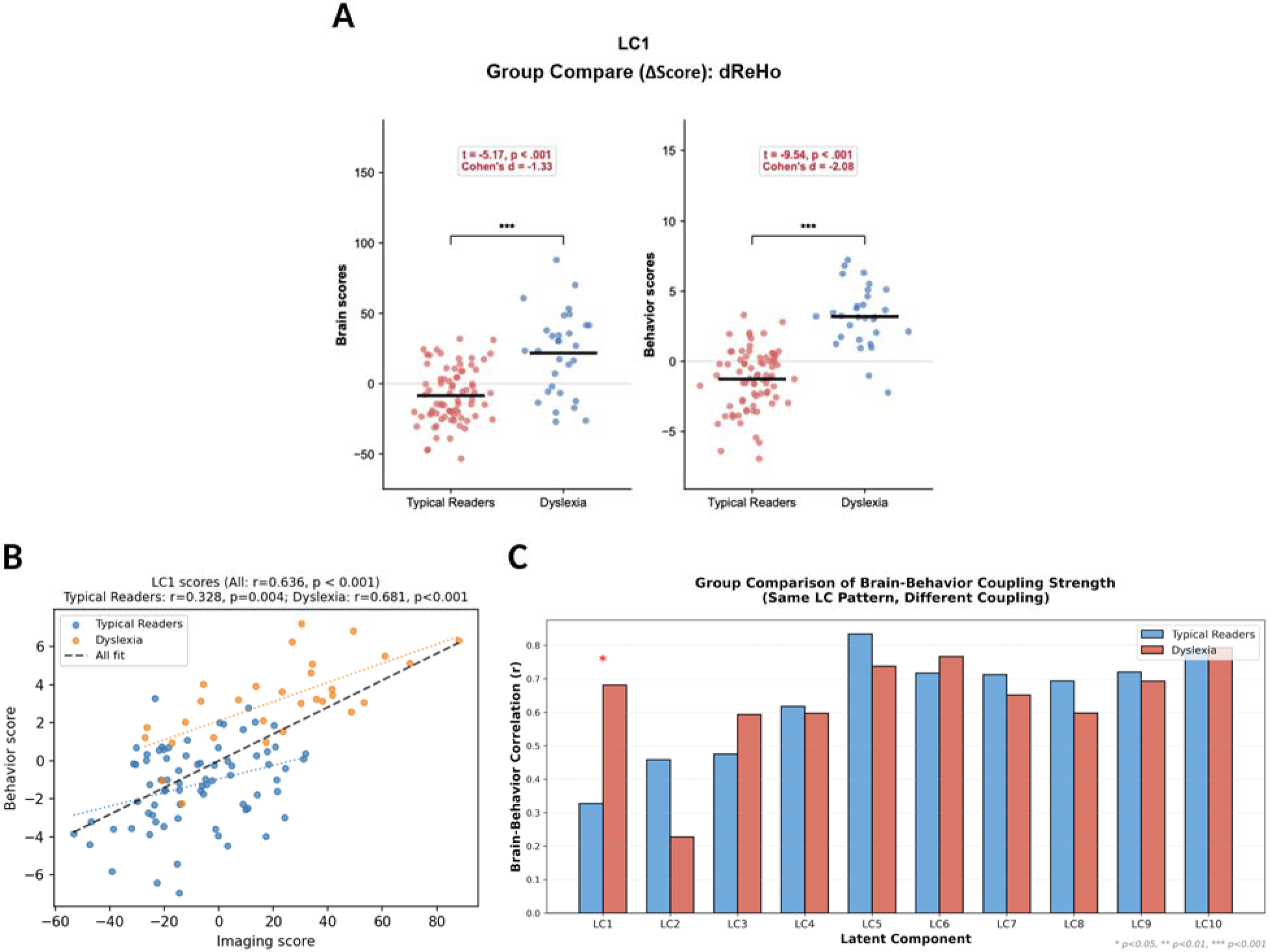
Group differences in brain-behavior associations revealed by Partial Least Squares Correlation (PLS-C) analysis of resting-state fMRI BOLD signal Dynamic Regional Homogeneity (dReHo) and the multi-domain behavioral model between typical readers (TR) and dyslexia (DY). (A) Group comparisons of LC1 brain and behavioral scores. Red: TR; Blue: DY. (B) Brain-behavior coupling strength (*r*) in LC1. Brain scores (x-axis) vs. behavioral scores (y-axis). Blue: TR; Orange: DY. (C) Brain-behavior coupling strength (Δr = rTR − rDY) across LC1-LC10. Significant differences mean brain-behavior coupling differed between groups on the same shared LC (\**p* < .05, \*\**p* < .01, \*\*\**p* < .001). Blue: TR; Red: DY. Statistical testing: Δr significance assessed via label permutation (2,000 iterations). TR: n = 76, DY: n = 30.

In the reading/language-specific model, LC1 did not reach significance in the group-comparison analysis (Figure S20A). Groups differed significantly in LC expression on brain (*t*(104) = −6.12, *d* = −1.39) and behavioral scores (*t*(104) = −8.49, *d* = −1.76; both *p* < .001; Figure S21A). Brain–behavior coupling did not differ between groups (*r*TR = 0.495 vs. *r*DY = 0.580; Δr = −0.085, *p*_permuted_ = .476; Figure S21B–C).

## Discussion

We used amplitude-based (SD, MSSD) and coupling-dynamics (dReHo) metrics as macroscale phenotypes relevant to NNH. First, rs-fMRI brain signal variability differed between adults with dyslexia and typical readers in a regionally specific rather than global manner.

Second, brain–behavior relationships differed between groups in a region- and metric-specific manner. Notably, dReHo uniquely differentiated adults with dyslexia from typical readers in brain–behavior coupling patterns across reading, spelling, math, and general cognition.

### Adult dyslexia is characterized by localized, metric-specific differences in brain signal variability

Voxel-wise analyses revealed region- and metric-specific effects: adults with dyslexia showed greater amplitude-based brain signal variability (SD and MSSD) in ventral occipitotemporal and cerebellar regions, whereas typical readers showed greater coupling-dynamics variability (dReHo) in dorsal temporoparietal and frontal regions (4,56). Control analyses (tSNR and meanFD) and whole-brain mean variability showed no group effects (Supplemental Results), reducing the likelihood that these effects reflect gross tSNR or head-motion differences, and suggesting region-specific rather than global signal differences.

SD/MSSD and dReHo were negatively correlated and showed opposite group effects (higher SD/MSSD in DY, higher dReHo in TR). Thus, greater brain signal variability is not uniformly “worse”; rather, its meaning may depend on metric and region (35,57). When restricted to reading-network masks, only dReHo showed group differences, suggesting that variability in coupling dynamics (dReHo) may be more proximal to disruption of reading-related functions (58), while amplitude-based metrics (SD/MSSD) may index instability across broader multi-domain functions.

### Brain-behavior covariance is region-, metric-, and domain-specific

Across TR and DY, we observed brain–behavior associations between amplitude-based brain signal variability (SD and MSSD) in visual, somatomotor, cerebellar, and subcortical networks and both multi-domain and reading/language-specific behaviors. These brain regions, which support reading automatization and timing (59), have previously been linked to multiple cognitive domains spanning phonological processing, executive control, and language efficiency (3,60), consistent with persistent deficits in reading fluency, pseudoword decoding, and spelling in adults with dyslexia despite prolonged literacy experience (61). Taken together, these findings suggest that the stability of moment-to-moment signal fluctuations in regions supporting automatization and timing relates to individual differences in both reading/language and broader cognitive skills, which may help explain why fluency, decoding, and spelling difficulties persist into adulthood in dyslexia. These continuous brain-behavior covariance patterns across adults with and without dyslexia also align with current theories that conceptualize psychiatric and neurodevelopmental conditions as variation along continua rather than discrete categories, such as the Research Domain Criteria (RDoC; 62), the Hierarchical Taxonomy of Psychopathology (HiTOP; 63), and transdiagnostic approaches to neurodevelopmental conditions (64). This conceptualization is consistent with the 2025 International Dyslexia Association (IDA) revised definition, which characterizes dyslexia as difficulties occurring along a continuum of severity (65), paralleling evidence from ADHD (66). Our findings therefore suggest that amplitude-based brain signal variability may index an individual’s position along a shared brain–behavioral continuum.

In contrast, local coupling-dynamics variability (dReHo) may reflect diagnosis-specific reorganization, as latent brain-behavior coupling with dReHo differed significantly between groups. Interestingly, however, this difference between adults with and without dyslexia was only observed in the multi-domain model (math, executive function, working memory, and reading) and not the reading/language-specific model, indicating cross-domain disruption of local dynamic coordination. This multivariate brain-behavior coupling pattern was also not dominated by canonical frontoparietal or salience networks, despite frontal regions appearing in voxelwise group contrasts. Together, these observations may suggest that dyslexia is characterized not only by differences in reading-related functions, but also by differences in more widespread relationships between local dynamics and broad multi-domain behaviors.

Overall, our findings highlight that dyslexia in adults may involve disruptions in multiple forms of brain signal variability, with amplitude-based metrics indexing continuous individual differences in reading-specific functions and metrics capturing temporal dynamics reflecting group-level reorganization of multi-domain functions.

### Limitations, Significance and Clinical Implications

Some limitations should be considered. Although rs-fMRI provides a task-free, clinically scalable measure of macroscale brain signal dynamics, it does not measure the cellular E/I mechanisms posited by the NNH. Moreover, BOLD brain signal variability may also reflect vascular, respiratory, and arousal-related fluctuations; future multimodal work (e.g., MRS, EEG, laminar fMRI) is needed to link these macroscale phenotypes to such mechanisms. PLS results should be interpreted with caution: cross-validation showed limited out-of-sample generalizability and spin tests attenuated significance, yielded near-zero out-of-sample correlations and SD/MSSD components did not survive spin tests, underscoring the need for replication in larger consortium samples (e.g., the ABCD Study, UK Biobank) across the lifespan (47). Spatial smoothing improves SNR but reduces local specificity; thus, dReHo differences should be interpreted as relative rather than absolute. The cross-sectional design also precludes causal inference, motivating longitudinal work.

Despite these limitations, the differing SD/MSSD and dReHo results speak to a key theoretical question in dyslexia: whether brain–behavior relationships vary continuously across readers (dimensional) or differ qualitatively between groups (categorical). Because group separation contributes to the behavioral covariance structure, group differences in LC expression alone cannot distinguish these accounts. Our results suggest that the answer depends on which aspect of brain dynamics is examined: amplitude variability appears to relate to behavior in the same way across readers, consistent with a continuum, whereas local coordination dynamics appear to relate to behavior differently in dyslexia, consistent with a categorical shift.

Dimensional and categorical views may thus be complementary descriptions of different levels of brain function rather than competing accounts. Moreover, because these brain–behavior relationships extended to math, working memory, and executive function, adult dyslexia may merit further investigation within multi-domain behavioral profiles rather than exclusively as an isolated reading deficit. Clinically, resting-state brain signal variability may offer a scalable task-free candidate neural phenotype of dyslexia (67) that could complement behavioral assessments, and our multi-domain findings may support further evaluation and support for adults with dyslexia beyond reading. Establishing out-of-sample generalizability of our findings in larger samples is a key next step toward individual-level application.

## Conclusions

Resting-state brain signal variability is not globally elevated across the brain in adults with dyslexia but differs from typical readers in a region- and metric-specific manner: amplitude variability relates to behavior along a shared continuum, whereas local coordination dynamics show group-dependent brain–behavior coupling spanning reading, math, working memory, and executive function. Pending replication in larger samples, these findings suggest that resting-state brain signal variability may be a promising scalable, task-free candidate brain phenotype of dyslexia capturing both reading-specific and multi-domain functions.

## Acknowledgments and Disclosures

### Author Contributions

J.L. contributed to conceptualization, investigation, data curation, methodology, formal analysis, visualization, project administration, writing–original draft, and writing–review and editing. A.S.K. contributed to writing–review and editing. Z.C.X. contributed to data curation, writing–review and editing. N.A. contributed to data curation, writing–review and editing. L.C.K. contributed to data curation, writing–review and editing. S.S.Y.C.L. contributed to writing–review and editing. K.P. contributed to writing–review and editing. F.H. contributed to supervision, funding acquisition, and writing–review and editing.

## Acknowledgment

We thank Sanhita Kancharla, an undergraduate research assistant, for assistance with figure preparation and manuscript proofreading.

## Data Availability

The data from this study are available upon reasonable request to J.L. and F.H.

